# Fed-batch production of antiviral „OP7 chimera” defective interfering influenza A particles

**DOI:** 10.64898/2026.09.30.755589

**Authors:** Wiebke Holste, Sascha Y. Kupke, Claudius Seitz, Udo Reichl, Holger Ziehr

## Abstract

Influenza A virus (IAV) defective interfering particles (DIPs) are regarded as promising broad-spectrum antivirals due to their high efficacy demonstrated in animal trials. Previously, an “OP7 chimera” DIP was generated that showed markedly enhanced antiviral efficacy compared to conventional IAV DIPs. In addition, a high-yield cell culture-based production process for OP7 chimera DIPs was established in perfusion mode. However, due to the technical complexity of this technology, we aimed to combine the advantages of perfusion in terms of yield with that of a simple and less complex batch process. In the present work, we established a fed-batch production process for OP7 chimera DIPs that allows a more straightforward implementation into existing production facilities. Using a feeding regime that was able to overcome the cell density effect, we achieved productive virus infection in stirred tank reactors until a cell density of 10E+6 cells/mL. Furthermore, we obtained a high virus titer of 3.8 log_10_(HAU/100 µL), corresponding to a 3.4-fold increase compared to a reference batch production. Moreover, the produced material demonstrated a significantly increased interfering bioactivity shown by an *in vitro* potency assay. Interestingly, this efficacy was similarly high as compared to OP7 chimera DIP material previously produced in perfusion mode, despite twofold lower virus titers. Finally, we demonstrate robustness of OP7 chimera DIP seed virus trains, which is significant step towards later pharmaceutical production. Further, corresponding GMP “MDCK-PB2(sus)” cell and “OP7 chimera” virus banks are now available for later biomanufacturing further use. In conclusion, the established fed-batch process offers a valid n additional option for pharmaceutical grade OP7 chimera DIP GMP manufacturing production under GMP conditions required for subsequent clinical testing and for later large-scale manufacturing.

**Key points:** - Robustness of OP7 chimera DIP seed virus production confirmed for GMP manufacturing

- Scalable fed-batch process enabled high infectious virus titers up to 7E+9 PFU/mL

- OP7 chimera DIP material harvested demonstrated very high antiviral activity

## Introduction

The influenza A virus (IAV) is a major human respiratory pathogen that poses a severe global health burden. Annual epidemics result in substantial morbidity and mortality while IAV pandemics, occurring every few decades, can trigger devastating societal and economic consequences and widespread human casualties (Krammer et al., 2018). The most effective preventive measure is annual vaccination (Tanner et al., 2021, Nypaver et al., 2021). In addition, due to their fast mode of action, small-molecule antivirals like oseltamivir or zanamivir can be used to complement vaccination programs during epidemics, and are additionally useful during pandemics (Jones et al., 2023, Holmes et al., 2021). However, the development of antiviral resistances of IAVs against such antivirals emphasizes the need for new antiviral treatment strategies.

One promising solution could be the use of virus-derived defective interfering particles (DIPs) (Smither et al., 2020, Levi et al., 2021, Li et al., 2021, Kalamvoki and Norris, 2022, Yin, 2025). Importantly, based on available data, the emergence of viral resistance to DIP-mediated inhibition appears unlikely (Chaturvedi et al., 2021, Dimmock and Easton, 2015, Pitchai et al., 2024, Rast et al., 2016). They occur within different viral species and have been shown to exert strong antiviral effects against corresponding wild-type (WT) viruses in different animal models (Dimmock et al., 2008, Welch et al., 2022, Rezelj et al., 2021, Zhao et al., 2022, Pitchai et al., 2024, Chaturvedi et al., 2021), as well as against unrelated virus species, conferring DIPs the potential for broad-spectrum antiviral use as well (Pelz et al., 2023, Lin et al., 2022, Xiao et al., 2021, Scott et al., 2011, Easton et al., 2011, Rand et al., 2021). IAV DIPs typically contain a large internal deletion in one of the eight viral RNA (vRNA) segments (Alnaji and Brooke, 2020, Dimmock and Easton, 2015), rendering them replication incompetent and therefore, potentially non-pathogenic. In a co-infection scenario with WT virus, the defective interfering (DI) genome (also termed defective viral genomes, delVGs or DVGs (Lohmann et al., 2025, Agu and Diaz-Munoz, 2025, Ranum et al., 2024, Sharov et al., 2021)) interferes with and suppresses WT genome replication, which constitutes an antiviral effect (Mendes and Russell, 2021, Rezelj et al., 2018, Vignuzzi and Lopez, 2019). In addition, DIPs stimulate antiviral interferon signaling, which adds up to the antiviral efficacy (Killip et al., 2015, Tapia et al., 2013, Yount et al., 2008, Vignuzzi and Lopez, 2019). As a result, IAV DIPs suppress many different respiratory viruses like IAV (Hein et al., 2021a, Hein et al., 2021c, Dogra et al., 2023, Dimmock et al., 2008), influenza B virus (Scott et al., 2011), respiratory syncytial virus (Pelz et al., 2023) and SARS-CoV-2 (Rand et al., 2021). IAV DIPs are intended to be administered nasally or orally using a droplet spray or nebulizer to treat the respiratory tract.

Previously, a new type of IAV DIP was discovered, termed “OP7”, that contains multiple point mutations in genome segment 7 (Seg 7-OP7), instead of a large internal deletion, as for conventional DIPs (cDIPs) (Kupke et al., 2019, Hein et al., 2021c, Rüdiger et al., 2024). Surprisingly, OP7 showed enhanced antiviral efficacy compared to cDIPs, as shown in *in vitro* and *in vivo* experiments (Hein et al., 2021a, Hein et al., 2021c, Dogra et al., 2023, Rand et al., 2021). In order to produce OP7 in bioreactors, an “OP7 chimera” DIP was generated using plasmid-based reverse genetics (Dogra et al., 2023). It contained the point mutations of the defective Seg 7-OP7 in addition to a large internal deletion in Seg 1. OP7 chimera DIPs were reconstituted in a mixture along with cDIPs, harboring a deletion in Seg 1 (Seg 1 cDIPs). Here, the WT Seg 7 of Seg 1 cDIPs can complement for the defect of OP7 chimera DIPs during a co-infection. In addition, complementing cell lines that expresses the “deleted” viral protein polymerase basic 2 (PB2), encoded on Seg 1, provide functional complementation for the Seg 1 defect. Together, it was shown that OP7 chimera DIPs can be produced along with Seg 1 cDIPs without infectious virus in complementing suspension Madin-Darby canine kidney (MDCK) cells, termed “MDCK-PB2(sus)”, in cell culture-based production processes (Dogra et al., 2023, Pelz et al., 2024).

So far, batch- and perfusion-mode processes were established for OP7 chimera DIP production in bioreactors (Pelz et al., 2024). Importantly, the fraction of OP7 chimera DIPs in relation to Seg 1 cDIPs can be controlled by the selection of an optimal multiplicity of infection (MOI) as well as certain process parameters, i.e. temperature and media dilution (MD) before infection (Dogra et al., 2023, Pelz et al., 2024). Doing so, a high fraction of approx. 99.9% of OP7 chimera DIPs could be achieved, which is important as it affects biological interfering efficacies (Pelz et al., 2024). Although a perfusion-based production allows for maximum yield, it would present challenges due to its increased complexity and the large volumes of medium that would need to be prepared and handled. Here, a fed-batch process would combine the advantages of both batch and perfusion production by enabling higher productivity with operational simplicity.

Therefore, in the present work, we established a high-titer fed-batch production process for OP7 chimera DIPs that could be swiftly implemented into existing standard production facilities. A high cell concentration of 10E+6 cells/mL could be achieved, using a fed-batch feeding profile that was capable to avoid the cell density effect (CDE). Compared to a previously published reference batch production process (Pelz et al., 2024), we achieved significantly higher virus titers and biological interfering efficacies of the produced material in fed-batch mode. Finally, even though virus titers were lower as compared to a previous perfusion process (Pelz et al., 2024), the interfering efficacies of the material produced in fed-batch mode were comparable, which demonstrate the potential of this process. Finally, “good manufacturing practice” (GMP) MDCK-PB2(sus) cell- and OP7 chimera DIP virus-banks were generated and are now available for further use. The easy adoptability of this process enables transfer into GMP production of OP7 chimera DIPs required for future clinical trials, which may pave the way for OP7 chimera DIPs as a novel antiviral broad-spectrum treatment modality.

## Materials and methods

### Cells and viruses

For IAV and OP7 chimera DIP production, complementing MDCK-PB2(sus) cells that had been genetically modified to express the viral PB2 protein (Hein et al., 2021a) were used. A research cell bank (RCB) was generated, which was then used for bioprocess development. MDCK-PB2(sus) cells were grown in chemically defined 4Cell CD MDCK medium (Sartorius), also known as Xeno-CD001S MDCK CD Medium (Bioengine), and 0.5 µg/mL puromycin in shake flasks (125 mL-1000 mL, baffled, with vent cap) at 37 °C, 5% CO_2_ and 180 rpm (50 mm orbit diameter). The adherent cell lines MDCK(adh) (ECACC, #84121903) and MDCK-PB2(adh), stably expressing PB2 (Bdeir et al., 2019), were used for virus quantification assays and were maintained in Glasgow’s minimal essential medium (GMEM) supplemented with 10% tryptose phosphate broth (TPB) and 10% fetal calf serum (FCS). For cell maintenance of MDCK-PB2(adh) cells, 1.5 µg/mL puromycin was added to the cell culture medium. Puromycin was only added during cell maintenance and not used in infection experiments. Viable cell concentration (VCC), viability and cell diameter were determined with a Cedex HiRes Analyzer (Roche).

Based on the OP7 chimera DIP seed virus (4.5E+6 PFU/mL) which was previously generated by reverse genetics and produced in MDCK-PB2(sus) cells, as described in (Dogra et al., 2023), a OP7 chimera DIP research virus bank (RVB, 1.3E+7 PFU/mL) was produced. This seed virus served both as the starting material for investigating the robustness of infection conditions (Fig. 1), for the manufacturing of GMP-compliant virus banks as well as for fed-batch production in a stirred tank bioreactor (STR) (Fig. 4). Influenza A/PR/8/34 (PR8, subtype H1N1), was used as standard virus (STV, 3.2E+8 TCID_50_/mL) in the interference assay, as well as reference in virus quantification assays. For fed-batch process development, a seed virus was produced from the STV in MDCK-PB2(sus) cells, hereafter referred to as IAV-PR8 (2.1E+9 PFU/mL).

**Figure 1:**
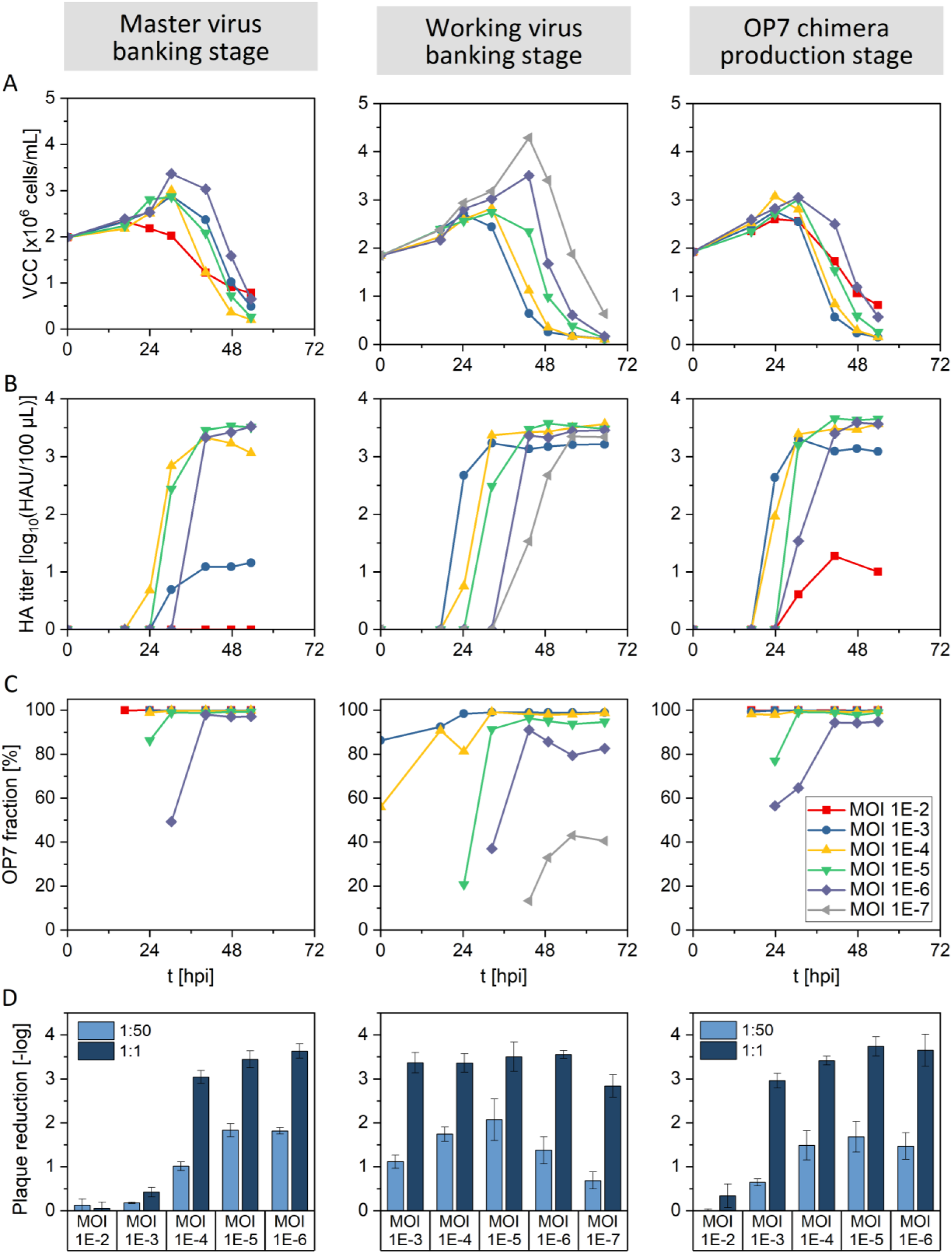
MOI-Screening for seed virus banking of OP7 chimera DIPs. MDCK-PB2(sus) cells were grown in shake flasks up to 4E+6 cells/mL, followed by infection with OP7 seed virus from the previous virus passage at different MOIs. Prior to infection a complete media exchange was performed, and cell density was adjusted to 2.0E+6 cells/mL. Infection conditions were as follows: 32 °C and 20 U/mL trypsin. (A) VCC. (B) HA titer. (C) OP7 fraction based on Seg 7 vRNA level, quantified by RT-qPCR. (D) Plaque titer reduction, determined in an interference assay. Parental MDCK(adh) cells were infected with STV (MOI 10, reference) or co-infected with OP7 chimera DIP material from optimal time of harvest (TOH). Plaque reduction was calculated using a plaque assay based on the reference plaque titer. Interference assay was performed in three independent experiments. Error bars indicate the standard deviation of three independent experiments. Dilutions of the tested materials are indicated. The TOH are listed hereafter from highest to lowest tested MOI: MVB “stage”: 32 hours post infection (hpi), 40 hpi, 40 hpi, 48 hpi, 48 hpi; WVB “stage”: 32 hpi, 43 hpi, 43 hpi, 49 hpi, 56 hpi; Production “stage”: 24 hpi, 31 hpi, 31 hpi, 41 hpi, 41 hpi.

### IAV-PR8 and OP7 chimera DIP production in shake flasks in batch mode

For seed virus banking of OP7 chimera DIPs and batch production, MDCK-PB2(sus) cells were seeded at a VCC of 0.5-0.7E+6 cells/mL in puromycin-free medium and grown in shake flasks to a VCC of 4E+6 cells/mL. For OP7 chimera banking and for investigating the robustness of infection conditions (Fig. 1), a complete medium exchange (CME) was performed prior to infection, in which the cell culture was centrifuged (200 g, room temperature, 5 min) and the cell pellet was resuspended in fresh medium to a VCC of 2E+6 cells/mL. For the batch reference process, a media dilution (MD) step was performed as described in (Pelz et al., 2024), in which cell culture was diluted 1:2 with pre-warmed fresh medium. Prior to infection 20 U/mL trypsin (Gibco) was added and cells were infected at MOI 1E-4, unless otherwise stated, and incubated at 32 °C, 5% CO_2_, 180 rpm. For sampling, samples of cell suspension were centrifuged and supernatant was stored in aliquots at -80 °C until further analysis.

For initial process development using IAV-PR8, MDCK-PB2(sus) cells were grown until the late exponential growth phase and infected either without, with MD, or with feed supplementation (Booster). For feed supplementation, 8%(v/v) HEK FS_2 (Sartorius) containing 33 mM glutamine was added prior to infection. The infection and sampling conditions were as described above.

### IAV-PR8 production shake flasks in fed-batch mode

For fed-batch production, MDCK-PB2(sus) cells were seeded at a VCC of 0.5 cells/mL and grown in shake flasks for five days, followed by infection. During the cell growth phase the cells were daily bolus-fed with HEK FS_2 (Sartorius) and GlutaMAX (Gibco) according to a fixed feeding regime based on off-line amino acid consumption data, as indicated. Prior to infection the last bolus feed was added and cells were infected at MOI 1E-4 using 20 U/mL trypsin (Gibco) and incubated at 32 °C, 5% CO_2_, 180 rpm as previously described for batch mode production.

### Bioreactor set-up, operation and infection

A 1L-STR (DASGIP parallel bioreactor system, Eppendorf) with a maximum working volume of 800 mL, equipped with two pitched-blade impeller (3-blade, 30° pitch, 50 mm diameter), and a L-sparger (macro-sparger), was used. Culture vessels were heated and cooled using a Bioblock (Eppendorf) connected to a Frigomix 1000 (Sartorius) recirculating cooler. The bioreactor was operated at 37 °C, pH 7.4 and 150 rpm. For virus replication phase, the temperature was reduced to 32 °C. Dissolved oxygen was maintained above 40% using a cascade control, in which air flow was increased first, before the oxygen in the gas flow was increased. The pH setpoint was controlled with CO_2_ gassing and addition of 7.5% (w/v) NaHCO_3_.

Prior to inoculation, MDCK-PB2(sus) cells, grown in shake flasks (as described before) were centrifuged (200 g, 5 min, room temperature) and then resuspended in fresh medium. The bioreactor was inoculated with a seed density of 0.5-0.7E+6 cells/mL, and the initial working volume (V_W_) was 400 mL (batch) and 650 mL (fed-batch). Feeding regime during cell growth phase for fed-batch mode were as described above, following a fixed feeding regimen, with the last feed bolus added immediately before infection. For batch mode production a MD step was carried out prior to infection. The V_W_ at time of infection (TOI) was approximately 750 mL. Infection and infection conditions were as described above.

### Extracellular metabolites

The concentration of glucose and lactate were quantified with a Stat Profile Prime CCS Analyzer (Nova Biomedical) or Biosen C-Line Analyzer (EKF Diagnostics). Ammonium concentration was measured using the Enzytec Liquid Ammonia kit (R-Biopharm). For quantification of amino acids a reverse-phase chromatography with an Agilent high-performance liquid chromatography (HPLC) 1200 system (Agilent Technologies) was used. A pre-column derivatization with ortho-phthalaldehyde (OPA) and 9-fluorenylmethylchloroformate (FMOC) was applied after which amino acids were separated using the ZORBAX Eclipse Plus C18 column (2.1 x 150 mm, 3.5 µm, Agilent) according to the Agilent ZORBAX Eclipse AAA protocol (Agilent).

The cell-specific uptake rates (q) were calculated for the respective metabolite in a given time interval as follows:

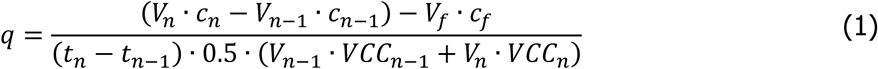

were V_n_ is the working volume at a given time (t_n_), c_n_ is the concentration of the respective metabolite at t_n_, V_f_ is the volume of added feed medium, c_f_ is the metabolite concentration in the feeding medium and VCC_n_ is the viable cell concentration at t_n_.

### Virus quantification

The total virus particle titer (tvp) was quantified by hemagglutination assay (HA assay) as described in (Kalbfuss et al., 2008). HA titer was expressed as log_10_ HA Units per 100 µL (log_10_(HAU/100 µL)). The total amount of IAV particles (virions/mL) was calculated as follows:

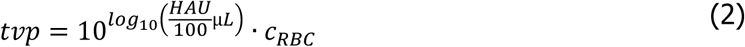

were c_RBC_ is the concentration of red blood cells (RBC) used in the HA assay.

Quantification of infectious virus titer was performed by plaque assay from (Hein et al., 2021a, Hein et al., 2021c). Infectious virus titer was expressed as plaque forming units per mL (PFU/mL). For quantification of OP7 chimera DIP seed virus titers, MDCK-PB2(adh) cells were used, allowing for detection of Seg1 cDIPs, which are propagation-competent in PB2-expressing cells and present in OP7 chimera DIP preparations. The relative standard deviation (RSD) was ≤41.8% based on independent measurements of the reference sample (n=29).

Virus productivity was evaluated based on cell-specific virus yield (CSVY) and volumetric virus productivity (VVP). CSVY and VVP were calculated based on total virus yield (vir_tot_) determined by HA assay or plaque assay as follows:

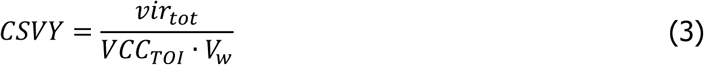

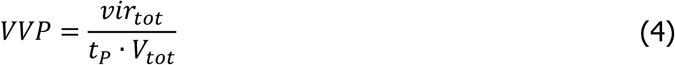

where VCC_TOI_ is the VCC at time of infection, V_w_ is the reactor working volume, t_P_ is the total process time and V_tot_ is the total volume of spent medium.

IAV vRNA segments were quantified by real-time RT-qPCR as described in (Kupke et al., 2019), using the Maxima H Minus Reverse Transcriptase (Thermofisher) for reverse transcription (RT) and the QuantiNova SYBR Green Mastermix (Qiagen) for qPCR. The primer sequences used for RT and qPCR to quantify wild-type Seg 7 (WT Seg 7) were derived from (Dogra et al., 2023), and for Seg 7-OP7 from (Hein et al., 2021c). For determining the relative proportion of OP7 chimera DIPs to Seg 1 cDIPs, the OP7 fraction was calculated based on Seg 7-vRNA levels as follows:

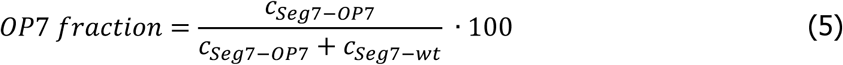

### Interference assay

Interfering efficacy of the produced OP7 chimera DIP material was evaluated *in vitro*, as previously described (Hein et al., 2021c). Briefly, MDCK(adh) cells were grown in 6-well plates and co-infected with 125 µL OP7 chimera DIP sample per well and with STV at an MOI of 10, or infected with STV only as a reference. In order to achieve a better resolution of the interfering effect, DIP samples were analysed both undiluted and 1:50 diluted. After incubation (37 °C, 5% CO_2_, 16 h), the supernatant was sampled and analysed by plaque assay, as described above. Plaque titer reduction was calculated, considering the plaque titer of the STV-only infection measurements.

### Host cell derived impurities

Double-stranded host cell DNA (dsDNA) was quantified with a Qubit 4 fluorometer (Thermofisher), using the Qubit 1X dsDNA HS Assay-Kit (Thermofisher). Host cell protein (HCP) concentration was measured with the MDCK HCP ELISA Kit (Cygnus).

## Results

### GMP cell and seed virus banking for production of “OP7 chimera” DIPs

The OP7 fraction is critical for the antiviral efficacy of produced OP7 chimera DIPs. However, fractions and virus titers were shown to be strongly dependent on the MOI used for production (Pelz et al., 2024, Dogra et al., 2023). With respect to seed virus banking, we investigated whether high fractions and titers can be maintained stably over multiple passages required for further GMP manufacturing.

For this, we first simulated the virus banking process under non-GMP conditions, using the MOI as the key control parameter. Based on previous work by (Dogra et al., 2023) a research virus bank (RVB) was produced based on a research cell bank (RCB), which both later served as biological starting material for the manufacturing of GMP cell and seed virus banks. To cover the necessary stages of future OP7 chimera production (Master Virus Bank (MVB), Working Virus Bank (WVB) and OP7 Production), two subsequent R&D seed virus banks were produced. MOI screenings were performed at each generated “stage” (Fig. 1), in order to determine optimal production conditions. The MDCK-PB2(sus) cells were grown in puromycin-free medium in shake flasks until a VCC of about 4E+6 cells/mL and a CME was performed in which cells were adjusted to 2E+6 cells/mL.

After infection, the cells continued to grow with maximum VCC being lowest and peaking faster for cells infected at higher MOIs compared to those infected at lower MOIs (Fig. 1A). In addition, higher MOIs (1E-2, 1E-3) resulted in lower total virus titers (Fig. 1B), while for lower MOIs the HA titers peaked later. Notably, the HA titer measured at MVB stage using a MOI of 1E-2 was even below the limit of detection, indicating a strong suppression of virus replication by OP7 chimera DIPs. In line with this, the proportion of OP7 chimera DIP, indicated as OP7 fraction based on the Seg 7-vRNA levels was higher towards higher MOIs (Fig. 1C). However, high OP7 fractions >99% could still be achieved up to MOI 1E-5, depending on the respective seed virus. Particularly unfavorable low OP7 fractions of only 43% were observed for production with MOI 1E-7, indicating an out-dilution of OP7 chimera DIP particles.

Next, antiviral efficacy was tested using an *in vitro* interference assay. For this, wildtype MDCK(adh) cells were co-infected with STV (MOI 10) and a fixed volume of the produced DIP samples. The STV suppression was then quantified using a plaque assay, in which plaque reduction was calculated based on a STV-only control. High interfering efficacy was observed when high HA titers were achieved in combination with a high OP7 fraction (Fig. 1D). Consequently, this resulted in an MOI optimum in the range of MOI 1E-4 to MOI 1E-6, in which a plaque reduction of 1.7 to 2.1 log was observed for 1:50 diluted DIP material. Based on these findings under R&D conditions, a GMP MVB and WVB were produced with the RVB serving as the starting material. For this, we used a GMP master cell bank (MCB), derived from the RCB. For production of the MVB and WVB, a MOI of 1E-5 and 1E-4 was used, respectively.

In summary, we show that higher MOIs can lead to inhibition of total virus titers, while towards lower MOIs, OP7 appears to get increasingly out-diluted. At intermediate MOIs, however, high OP7 fractions and virus titers can be maintained over a large range of MOIs (10E-4 to 10E-6), indicating robustness of OP7 chimera propagation over several passages, relevant for seed virus trains for GMP production. Finally, the GMP MDCK-PB2(sus) cell and OP7 chimera DIP virus banks are now available for further use.

### High-yield fed-batch production of IAV in shake flasks

A complete media exchange prior infection (Fig. 1) is not possible at larger scales without cell retention devices. In order to increase yields in a batch process, a media dilution (MD) step (1:2 dilution) can be implemented by adding fresh media (Pelz et al., 2024), but this results in low cell concentrations. To further intensify the process, a higher cell density is required while avoiding the cell density effect. Therefore, we established a fed-batch production process for IAV in shake flasks (Fig. 2). Here, the backbone virus for OP7 chimera DIPs (IAV-PR8) was used for process development.

**Figure 2:**
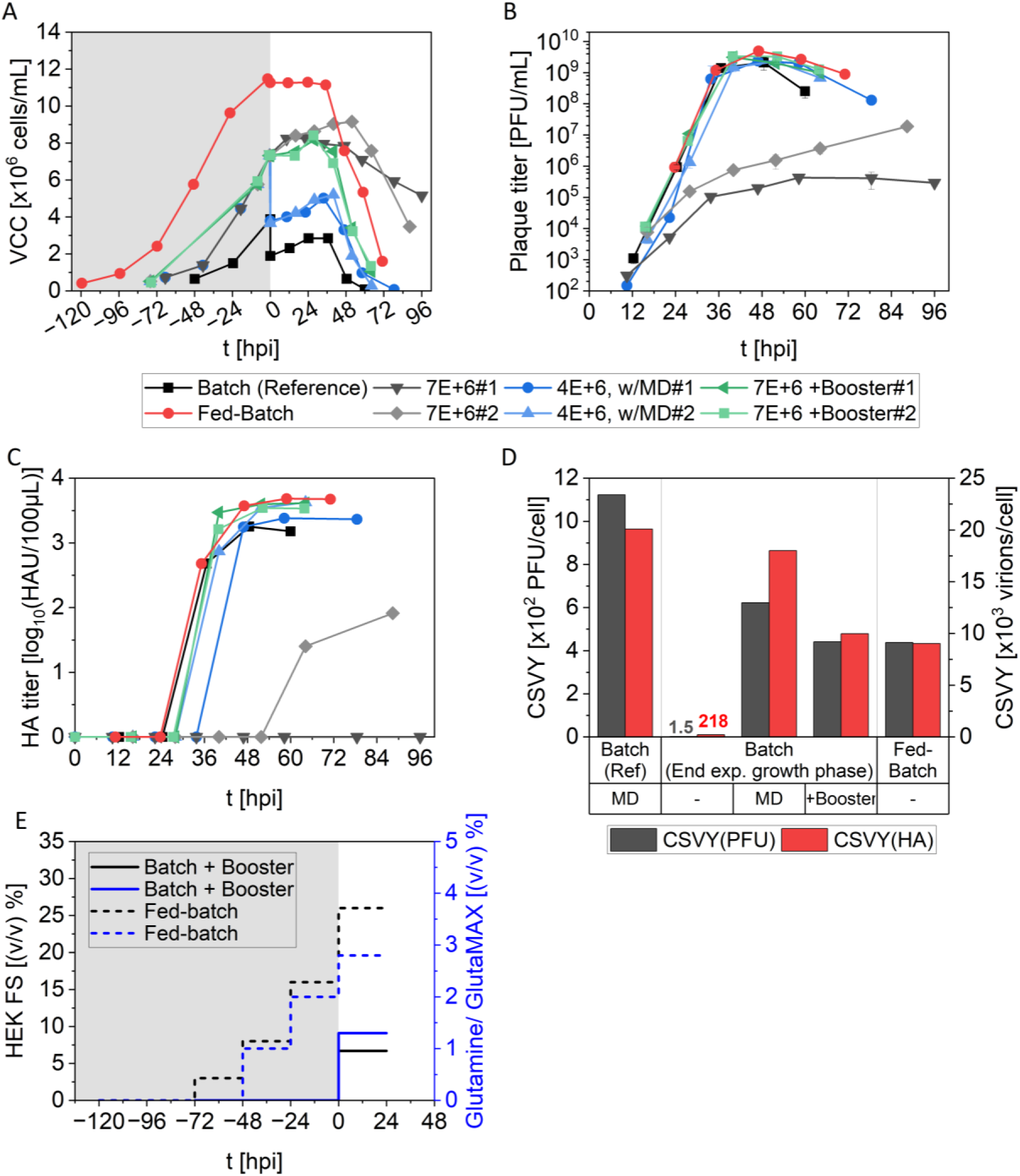
Establishment of a fed-batch process in shake flasks for IAV-PR8 production. MDCK-PB2(sus) cells were seeded between 0.5 E+6 to 0.7E+6 cells/mL and grown in shake flasks. Cells were cultured in batch mode for either 2 days (batch reference) or 3 days (end of exponential growth phase), or in fed-batch mode for 5 days, followed by infection with IAV-PR8. For fed-batch production, the cells were daily bolus-fed with HEK FS_2 and GlutaMAX. Prior to infection, a media dilution step (MD) was performed for the batch reference production, in which the cell suspension was diluted 1:2 with fresh medium. For cells grown in batch mode until late exponential growth phase, infection without medium or feed addition (7E+6), infection with MD step (7E+6, w/MD), and infection with feed supplementation (7E+6 +Booster) were tested. For the latter, cells were supplemented with 8%(v/v) HEK FS_2 containing 33 mM glutamine prior to infection. Infection conditions were as follows: 32 °C, MOI 1E-4 and 20 U/mL trypsin. Infection experiments in batch mode at the end of exponential growth phase were performed in two independent replicates. (A) VCC. (B). Plaque titer. Error bars show standard deviation of three independent plaque assays. (C) HA titer. (D) Cell specific virus yield (CSVY), based on plaque and HA titer. (E) Feeding regime.

In order to characterize IAV-PR8 production at increased cell density, three infection strategies were initially tested at increased cell density in batch mode: infection without medium or feed addition (7E+6 cells/mL), infection with MD step (7E+6, w/MD), and infection with feed supplementation (7E+6 +Booster). For this, MDCK-PB2(sus) cells were grown in shake flasks until late exponential growth phase (about 7E+6 cells/mL) and either infected directly (7E+6), diluted at a ratio of 1:2 with fresh medium (7E+6, w/MD), or 8% HEK FS_2 feed medium supplemented with 33 mM glutamine was added prior to infection (7E+6 +Booster). In addition, the previously developed batch process (Pelz et al., 2024) was performed as a reference, in which cells were grown until about 4E+6 cells/mL and infected after a MD step. For all conditions, the temperature was lowered to 32 °C prior to infection, followed by the addition of 20 U/mL trypsin and infection of the cells at an MOI of 1E-4.

Infection at increased cell density resulted in reduced cell specific virus yields (CSVY), when compared to the batch reference process (Fig. 2D). For infection at elevated VCC, both the MD step and feed supplementation prior to infection improved CSVY(PFU) by 408-fold and 288-fold, compared to the production without MD or Booster. The latter resulted in a highly unproductive state with delayed virus production and maximum plaque titers ranging from 4.3E+5 PFU/mL to 1.9E+7 PFU/mL, and maximum HA titers ranging from below the limit of detection to 1.9 log_10_(HAU/100 µL). In contrast, feed supplementation led to a significant increase in virus titers up to 3.3E+9 PFU/mL (Fig. 2B) and 3.60 log_10_(HAU/100 µL) (Fig. 2C) compared to the batch reference (2.1E+9 PFU/mL, 3.25 log_10_(HAU/100 µL). In line with the high titers observed, the amino acid concentrations before and after feed supplementation showed that all consumed amino acids had been replenished to a level comparable with the batch reference (Fig. S1).

Next, a fixed feeding regime was calculated for fed-batch production based on previous amino acid consumption data (not shown), using the HEK FS_2 feed and GlutaMAX (Fig. 2E). This aimed to replenish essential nutrients at time of infection (TOI), using serine, valine, isoleucine, leucine and arginine as a feed indicator. A previous dynamic feeding regime based on at-line glucose concentration measurements had not been successful in overcoming the CDE at higher VCC (data not shown). Therefore, cells were seeded at 0.5E+6 cells/mL and grown for five days until about 11E+6 cells/mL, followed by infection. As shown in Fig. 2D, cells were daily bolus-fed starting after 2 days post inoculation, whereas GlutaMAX was only added in low levels, since the used 4Cell MDCK CD medium is already supplemented with a high glutamine concentration of 8 mM by the manufacturer. The fed-batch process allowed the CSYV to be maintained at a medium level of 438 PFU/cell (Fig. 2D), which was lower than in the batch reference process. However, the fed-batch approach enabled infection at a significantly higher VCC. As a result, the virus titer increased up to 4.9E+9 PFU/mL (Fig. 2B) and 3.7 log_10_(HAU/100 µL) (Fig. 2C), while virus titer dynamics were comparable with the batch reference process, with maximum virus titers reached after about 48 hours post infection (hpi). Moreover, no nutrient limitations were observed for the fed-batch production process (data not shown).

Taken together, we established a high cell density and high-yield fed-batch production of IAVs in shake flasks with up to 11E+6 cells/mL, where the feeding regime allowed overcoming the cell density effect. Compared to the reference batch production process, we achieved an increase in infectious and total virus titers of 2.3-fold and 2.7-fold, respectively.

### Fed-batch production of IAV in a bioreactor

To investigate the scalability of the established fed-batch production, we transferred the process from the shake flask to a stirred tank reactor (STR). Next, we compared the fed-batch production (FB) to the reference batch process (B) in STRs(Pelz et al., 2024) and to corresponding productions in shake flasks (SF). For this, MDCK-PB2(sus) cells were seeded at 0.5E+6 cells/mL (FB-STR#1) and 0.7E+6 cells/mL (FB-STR#2, B-STR) and cultivated until a VCC of about 4E+6 cells/mL (batch) or 10E+6 cells/mL (fed-batch), followed by infection. While for FB-STR#1, the cells were daily bolus-fed during the cell growth phase with HEK FS_2 and GlutaMAX according to the shake flask production (FB-SF), feed addition for FB-STR#2 was adjusted in accordance with the slightly higher seed density (Fig. 3G). Prior to infection, either a final bolus feed was added (fed-batch) or an MD step was performed (batch), resulting in a final working volume of approximately 750 mL. As in the shake flasks, the temperature was shifted to 32 °C, 20 U/mL trypsin was added, and the cells were infected at a MOI of 1E-4.

**Figure 3:**
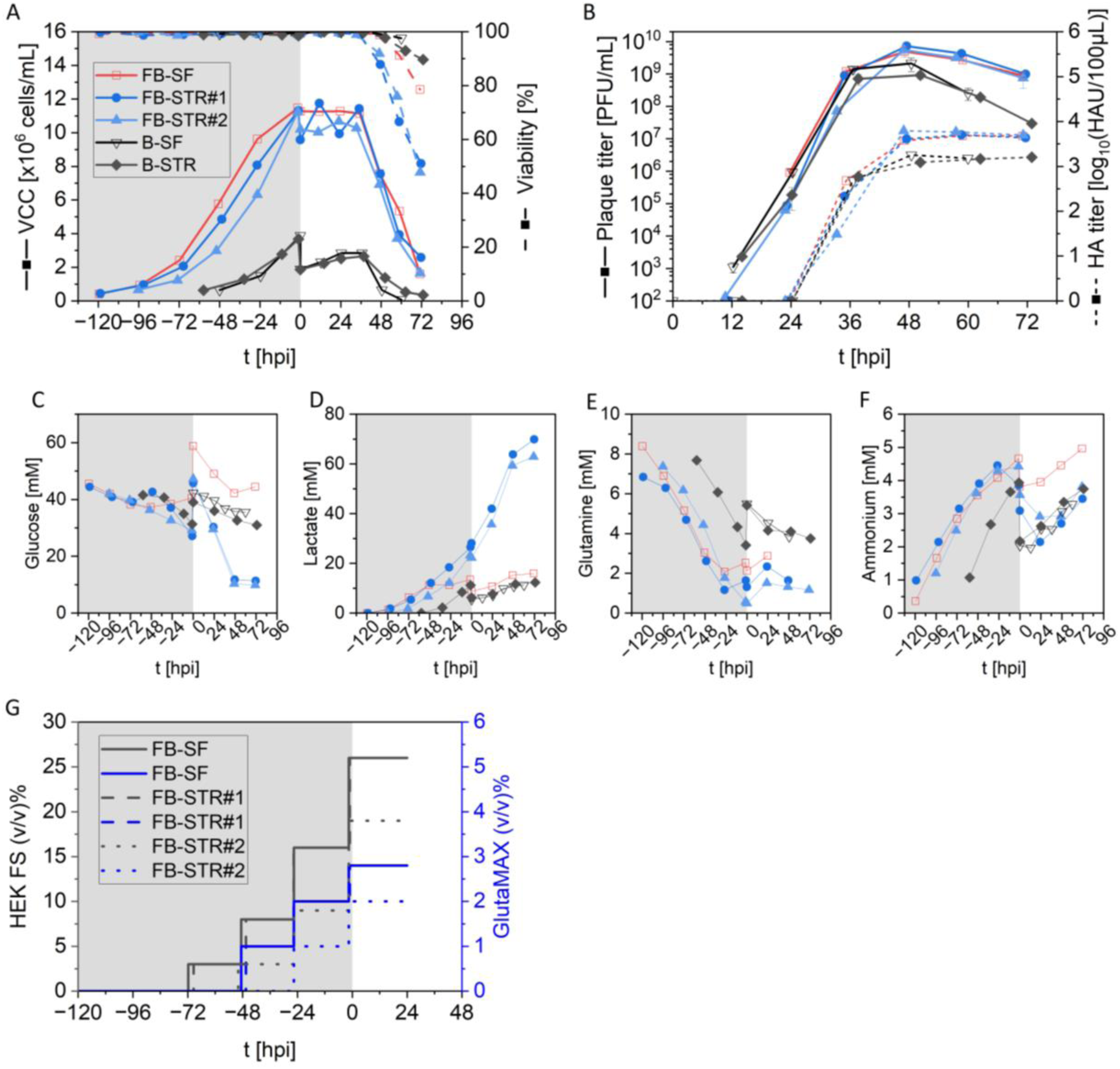
Comparison of IAV-PR8 production in in fed-batch mode and batch mode in a 1L-STR compared to shake flask production (SF). MDCK-PB2(sus) cells were seeded between 0.5E+6 to 0.7E+6 cells/mL and grown in batch or fed-batch-mode to a VCC of 4E+6 cells/mL or 10E+6 cells/mL, respectively. For fed-batch production (FB-SF, FB-STR), cells were daily bolus fed with HEK FS and GlutaMAX. For batch production (B-SF, B-STR), a media dilution step was performed prior to infection. Infection conditions were as follows: 32 °C, MOI 1E-4 and 20 U/mL trypsin. (A) VCC (solid line) and viability (dashed line). (B) Plaque titer (solid line) and HA titer (dashed line). Error bars show standard deviation of three independent plaque assays. (C-F) Main metabolite concentration. (G) Feeding regime. Note that data for shake flask production (open symbols) are taken from figure 1.

Despite the differences in production scale and process control, very comparable results were achieved between shake flask and STR productions with regard to both cell growth (Fig. 3A) and virus titer dynamics (Fig. 3B). Only minor differences were observed in the fed-batch production during the cell growth phase, with cells in the STR showing slightly lower growth rates than in the shake flask. The STR infections in fed-batch mode yielded maximum plaque titers of 7.2E+9 PFU/mL (FB-STR#1) and 5.5E+9 PFU/mL (FB-STR#2), which was slightly higher than the previous shake flask infection (4.9E+9 PFU/mL). In contrast, the batch STR run resulted in a slightly reduced plaque titer of 9.2E+8 PFU/mL compared to batch mode production in shake flask (2.1E+9 PFU/mL). With regard to the volumetric virus productivity (VVP) based on plaque titer, the fed-batch process resulted in a 4.8-fold improvement compared to the batch reference process (Tab. 1).

**Table 1:** Overview of process performance of IAV-PR8 and OP7 chimera DIP production in batch and fed-batch) mode in shake flask and 1L-STR.

|  | B-SF | FB-SF | B-STR#1 | B-STR#2 | FB-STR#1 | FB-STR#2 | FB-STR#3 |
| --- | --- | --- | --- | --- | --- | --- | --- |
| IAV | PR8 | PR8 | PR8 | OP7 chimera | PR8 | PR8 | OP7 chimera |
| VCC at TOI [cells/mL] | 1.9E+6 | 11.3E+6 | 1.9E+6 | 2.1E+6 | 9.6E+6 | 10.2E+6 | 10.0E+6 |
| TOH (PFU) [hpi]<sup>a</sup> | 49 | 47 | 50 | 39 | 48 | 47 | 46 |
| Max. plaque titer [PFU/mL] | 2.1E+9 | 4.9E+9 | 9.2E+8 | n/a | 7.2E+9 | 5.5E+9 | n/a |
| CSVY [PFU/cell]<sup>b</sup> | 1123 | 438 | 492 | n/a | 751 | 536 | n/a |
| VVP [PFU/L/d] | 5.3E+11 | 6.2E+11 | 2.0E+11 | n/a | 1.0E+12 | 9.1E+11 | n/a |
| HA titer [log<sub>10</sub>(HAU/100µL)] | 3.25 | 3.57 | 3.09 | 3.21 | 3.61 | 3.79 | 3.80 |
| Total virus titer [virions/mL] | 3.8E+10 | 7.9E+10 | 2.5E+10 | 3.3E+10 | 8.1E+10 | 1.2E+11 | 1.3E+11 |
| CSVY<sub>HA</sub> [virions/cell]<sup>b</sup> | 2.0E+4 | 7.0E+3 | 1.3E+4 | 1.6E+4 | 8.4E+3 | 1.2E+4 | 1.3E+4 |
| VVP<sub>HA</sub> [virions/L/d] | 9.4E+12 | 1.1E+13 | 5.5E+12 | 8.1E+12 | 1.2E+13 | 2.1E+13 | 2.1E+13 |
| Seg 7-OP7 vRNA [vRNA/mL] | n/a | n/a | n/a | 7.7E+10 | n/a | n/a | 1.4E+12 |
| OP7 fraction [%]<sup>c</sup> | n/a | n/a | n/a | 98.5 | n/a | n/a | 99.6 |
| dsDNA [µg/mL] | - | - | 12.3 | 3.2 | 39.0 | 37.4 | 30.8 |
| HCP [µg/mL] | - | - | 70.0 | 8.9 | 257.6 | 303.3 | 202.3 |

Note: B, batch; CSVY, cell specific virus yield; FB, fed-batch; SF, shake flask; TOH, time of harvest; VVP, volumetric virus productivity
<sup>a</sup>TOH was based on max. plaque titer (IAV-PR8) or max. interfering efficacy (OP7 chimera DIP)
<sup>b</sup>CSVY was calculated based on VCC at TOI
<sup>c</sup>OP7 fraction was calculated based on vRNA levels

Although good comparability was observed for fed-batch mode production between shake flask and STR cultivation, clear differences in metabolite consumption and production became evident (Fig.3C-F). For the FB-STR cultures, the glucose (Fig. 3C) and amino acid data (Fig. S2) indicate notably higher consumption rates, compared to the shake flask culture. However, as glucose and amino acid concentration did not indicate nutrient limitations, this does not seem to compromise the fed-batch process in the STR. Interestingly, higher lactate formation was accompanied with decreased ammonium concentration in the STR scale for fed-batch production. Lactate concentrations in the FB-STRs reached levels of up to 64 mM, which was 4-times higher than in the shake flask culture and 5-times higher than in the batch reference STR (Fig. 3D). In contrast, ammonium levels in the FB-STRs started to decrease at the end of cell growth phase, resulting in ammonium concentrations of only 2.9 mM at 48 hpi, whereas in the shake flask the amomum level increased up to 4.5 mM (Fig. 3E). The ammonium concentration in the batch-reference STR was 3.3 mM. Overall, the metabolite concentrations and virus titers indicate that there were no limiting or inhibitory concentrations in either batch or fed-batch production.

In summary, the process transfer of fed-batch production from a shake flask to a STR resulted in comparable virus titers. In addition, fed-batch productions in STRs were highly reproducible. Compared to the reference batch production process in STRs, the fed-batch productions yielded a 6.9-fold and 3.4-fold increase in infectious and total virus titers, respectively. Together, our data indicates that further scale-up to industrial volumes appears straightforward to accomplish.

### High-yield fed-batch production of “OP7 chimera” DIPs in a bioreactor

In order to finally validate the established fed-batch process for production of the OP7 chimera, we compared the fed-batch process with the batch reference process in STRs with regard to OP7 chimera DIP production (Fig. 4). Therefore, MDCK-PB2(sus) cells were grown in fed-batch mode and batch mode, followed by infection after 96 h (fed-batch) and 58 h (batch), as described above. For infection, the previously produced OP7 chimera seed virus (Fig. 1, “production stage”) was used instead of the IAV-PR8 backbone virus, using an MOI of 1E-4.

**Figure 4:**
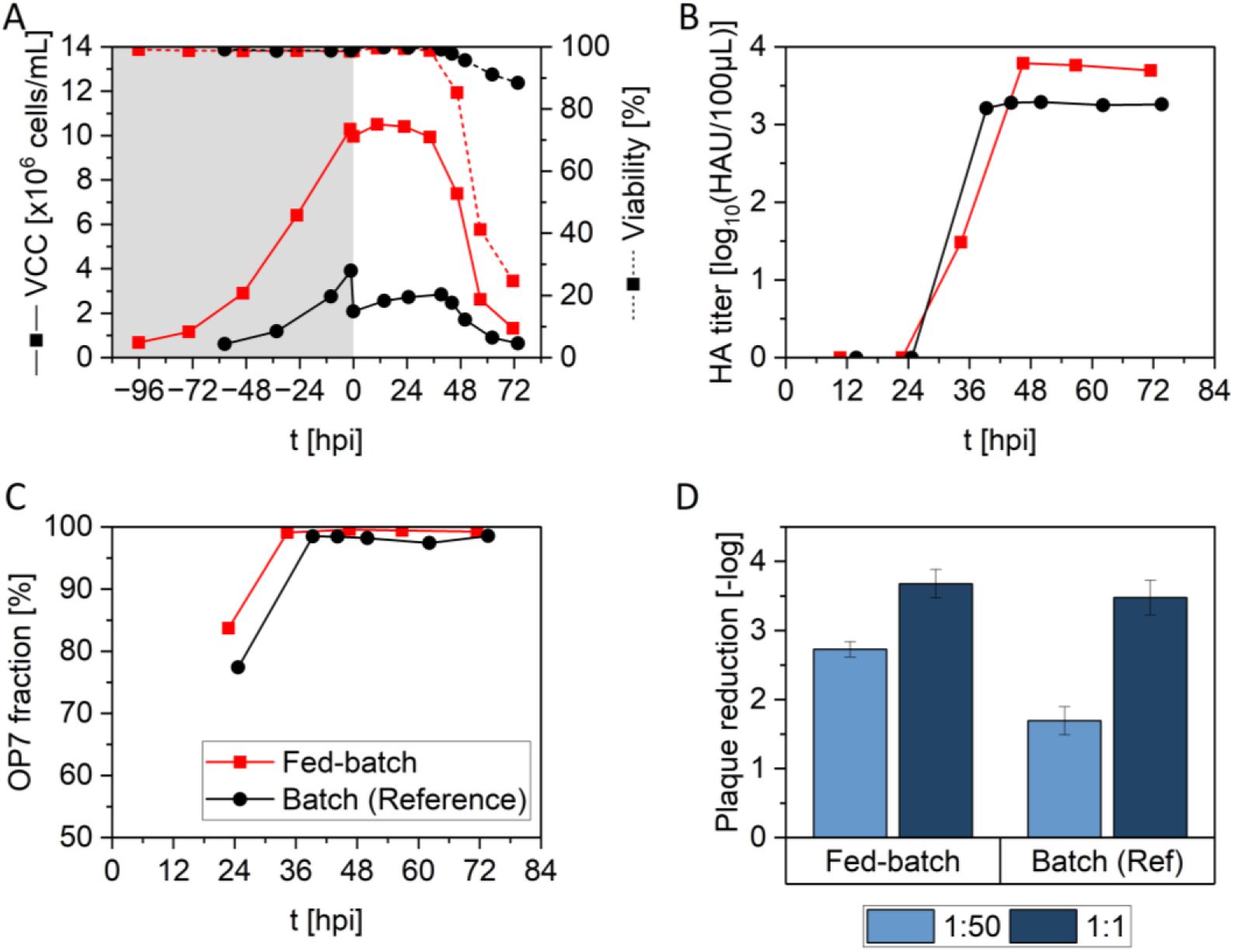
OP7 chimera production in fed-batch mode in a 1L-STR versus the batch reference process. MDCK-PB2(sus) cells were infected with OP7 chimera DIPs from working virus banking stage (Fig. 1). For fed-batch production, cells were daily bolus fed with HEK FS and GlutaMAX (like FB-STR#2, Fig. 3G). For the batch reference process, a media dilution step was performed prior to infection, in which the cell suspension was diluted 1:2 with fresh medium, according to Pelz et al. (2024). Infection conditions were as follows: 32 °C, MOI 1E-4 and 20 U/mL trypsin. (A) VCC. (B) HA titer. (C) OP7 fraction, based on Seg 7-OP7 and WT Seg 7 vRNA levels, quantified by RT-qPCR. (D) Interference assay. For evaluation of *in vitro* interfering efficacy, MDCK(adh) cells were infected with STV (MOI 10) alone (reference) or co-infected with 125 µL OP7 chimera DIP material, after which plaque reduction was quantified by plaque assay based on reference infection. Error bars indicate the standard deviation of three independent experiments. Dilutions of the tested materials are indicated. The optimal TOH: 46 hpi (fed-batch), 39 hpi (batch).

The cell growth and virus titer dynamics were comparable to that of the previous IAV-PR8 STR runs, as indicated by the growth curve (Fig. 4A) and HA titer results (Fig. 4B). After infection, the VCC remained constant until 34-39 hpi, after which cell lysis started. The total virus titer reached its maximum value of 3.8 log_10_(HAU/100 µL) after 46 hpi (fed-batch), and 3.3 log_10_(HAU/100 µL) after 44 hpi (batch). In addition, high OP7 fractions of 99.6 % (fed-batch) and 98.5% (batch) were reached (Fig. 4C), as expected for the applied production MOI (Fig. 1). Next, the produced OP7 chimera DIP material was tested for antiviral activity using the *in vitro* interference assay (Fig.4D) with samples harvested at the respective TOH (fed-batch: 46 hpi, batch: 39 hpi). The ideal TOH coincided with reaching the HA plateau and the onset of VCC decline due to virus-induced cell lysis, and avoid cell debris that could impair the subsequent DSP process and thus the overall yield. The OP7 chimera material produced in fed-batch mode exhibited significantly higher interfering activity, with a reduction in the release of infectious STV particles of 2.73 log steps for the diluted samples (1:50). In comparison, the batch reference material resulted only in a reduction of 1.60 log steps.

As residual host cell derived DNA and host cell derived proteins (HCP) in biologicals can be a serious safety risk, there are regulatory requirements regarding their characterization and reduction. Since impurities are reduced during purification in the downstream processing (DSP) part, we characterized dsDNA and HCP level from the STR runs at optimal TOH, to provide more comprehensive data as starting point for establishment of a future DSP process. As for the fed-batch process, cells were infected at higher VCC, which also resulted in higher dsDNA and HCP level (Tab. 1). However, the ratio of impurity per virus particle was very comparable. Table 1 summarizes the process performance including virus titer, CSVY, VVP and host cell derived impurities for both IAV-PR8 and OP7 chimera production. The results from three independent FB-STR runs demonstrates good reproducibility regarding process productivity, including process time and virus titer and impurity level.

Taken together, we verify the established fed-batch process in STRs for production of OP7 chimera DIPs. Compared to the reference batch production process, we achieved a 3.4-fold higher total virus particle yield. In addition, a comparably high purity of 99.6% of OP7 chimera DIPs was achieved along with a significantly increased interfering efficacy of the produced OP7 chimera DIP material compared to the batch production process.

## Discussion

In the present study, we established a technically scalable and industry-compatible fed-batch production process for antiviral “OP7 chimera” DIPs. Given the technical complexity associated with previously established perfusion-based processes, the development of a robust fed-batch strategy represents an important step towards simplified GMP-compliant manufacturing. Here, we demonstrate that high cell density cultivation in fed-batch mode effectively mitigated the cell density effect and enabled substantially increased virus yields compared to conventional batch production. Notably, despite lower overall virus titers than previously reported for perfusion cultivation, the antiviral efficacy of fed-batch-derived material was comparable. Furthermore, robustness of OP7 chimera propagation across multiple passages was demonstrated, confirming the suitability of the established cell and seed virus banks for GMP manufacturing.

Our data demonstrate a pronounced MOI dependency of OP7 chimera DIP production, confirming previous studies (Dogra et al., 2023, Pelz et al., 2024), indicating that balanced infection conditions are essential for simultaneously achieving both, high virus titers and high OP7 fractions. While high MOIs resulted in elevated fractions of OP7 chimera DIPs, they were also associated with reduced total virus yields. This suggests an out-growth of OP7 chimera DIPs compared to Seg 1 cDIPs, impairing overall virus propagation (Dogra et al., 2023). In contrast, lower MOIs tendentially promoted higher virus titers, likely due to increased propagation of Seg 1 cDIPs at the expense of OP7 chimera DIPs, which are out-diluted. Intermediate MOIs, however, enabled maintenance of both high OP7 fractions and high virus titers. This behavior can be explained by the unique biological dependency of OP7 chimera DIPs on co-infection with Seg 1 cDIPs (Dogra et al., 2023, Pelz et al., 2024). Since OP7 chimera DIPs are replication-deficient, even in PB2-complementing cells, efficient propagation requires simultaneous presence of Seg 1 cDIPs providing functional WT Seg 7 vRNA. At high MOIs, co-infection events occur more frequently, thereby promoting complementation and preferential amplification of OP7 chimera DIPs, which outcompete Seg 1 cDIPs. Consequently, propagation of the complementing Seg 1 cDIPs becomes increasingly suppressed, ultimately limiting total virus production. Conversely, under low-MOI conditions, single-hit infections predominate, favoring Seg 1 cDIP replication while propagation-deficient OP7 chimera DIPs cannot replicate and become low-abundant in the virus population.

Importantly, however, our data additionally demonstrate a comparatively broad operational window in which favorable production characteristics can be maintained, with an optimal MOI range around 10E-4 to 10E-6. This robustness is particularly relevant in the context of GMP manufacturing, where reproducible process performance parameters and consistent product quality attributes are essential prerequisites. In this regard, the MOI represents a, if not even the most, critical process parameter (CPP), as it directly influences the biological composition of the produced material and thereby its antiviral activity, which is considered a critical quality attribute (CQA), which both will play important roles in later process validation. The broad acceptable MOI range substantially reduces the likelihood of operating under unfavorable infection conditions and therefore lowers the risk of inconsistent product quality or costly batch failure during manufacturing.

Compared to established batch and perfusion processes, fed-batch cultivation represents an attractive compromise between process simplicity and production performance for virus manufacturing. Conventional batch processes are characterized by ease of operation, low technical complexity, and robust process handling, but are inherently limited by cell densities achieved and therefore restricted virus yields (Silva et al., 2021, Pelz et al., 2024, Hein et al., 2023, Kuchler et al., 2022). In contrast, perfusion processes enable substantial process intensification, but typically achieve very high productivities and space-time yield (STY) for virus production (Silva et al., 2023, Pelz et al., 2024, Wu et al., 2021, Zinnecker et al., 2025). However, these advantages come at the expense of considerably increased process complexity, including the requirement for high perfusion rates, large media consumption, additional equipment, membrane fouling risks, and elevated shear stress (Hein et al., 2021b, Silva et al., 2023, Wu et al., 2021). Fed-batch cultivation, by comparison, is currently the most frequently applied bioprocess mode in the biopharmaceutical industry due to its operational simplicity and scalability (Chen et al., 2018, Matanguihan and Wu, 2022). Nevertheless, its potential for influenza A virus production has so far received only little attention, and only few examples have been reported in the literature (Granicher et al., 2019, Silva et al., 2023, Vazquez-Ramirez et al., 2019). Notably, these reported approaches represent hybrid concepts that combine fed-batch cultivation during the cell growth phase with perfusion during virus replication.

Although the chemically defined 4Cell MDCK medium is well established for influenza virus production in MDCK cells in both batch and perfusion mode (Hein et al., 2021b, Hein et al., 2021a, Hein et al., 2021c, Pelz et al., 2024, Wu et al., 2021, Bissinger et al., 2021), its suitability for IAV DIP production had not yet been demonstrated. In the present study, a major challenge during process development was the occurrence of the CDE, where increasing VCCs resulted in reduced CSVYs, which was observed in the batch cultivations and fed-batch cultivation using an unsuitable feeding regime. Similar observations have previously been described for IAV and adenovirus production in HEK cells (Silva et al., 2023, Joe et al., 2022, Shen et al., 2024).

While the exact cause of the CDE remains unclear, both accumulation of inhibitory compounds and changes in cell physiological state have been proposed as contributing factors (Lavado-Garcia et al., 2022). In many virus production processes, the CDE can be alleviated by media exchange or perfusion-based feeding strategies that replenish nutrients while simultaneously diluting inhibitory metabolites (Henry et al., 2004, Dill et al., 2019, Shen et al., 2024). In the present study, both the implementation of a media dilution step and the feed supplementation using the commercial HEK FS feed restored virus titers at elevated cell concentrations.

Further optimization of the fed-batch process demonstrated that nutrient availability, in particular amino acid supply, played a central role in maintaining productive virus replication at elevated cell densities. Amino acid analyses revealed potential nutrient limitations at TOI, prompting refinement of the feeding regime based on offline amino acid monitoring. This adapted strategy maintained CSVYs at productive levels, indicating that amino acid availability represented a suitable indicator for feed supplementation using the HEK FS medium. Nevertheless, nutrient supply alone is unlikely to fully explain process performance, as accumulation of metabolites such as lactate and ammonium has also been associated with impaired virus production (Genzel et al., 2005, Kegel et al., 2025). In the present fed-batch shake flask cultivations, ammonium concentrations reached up to 4.5 mM during productive infection, but these levels did not negatively affect virus yields, despite previous reports linking elevated ammonium concentrations to reduced virus productivity (Kegel et al., 2025, Ferreira et al., 2007, Salvi et al., 2002).

To evaluate process scalability, the established fed-batch strategy was subsequently transferred from SFs to STRs, enabling controlled cultivation conditions including regulated pH and oxygen supply. The process transfer was successful and resulted in robust virus production with high titers comparable or superior to those achieved in SF cultivations. Notably, fed-batch production in stirred tank reactors yielded higher virus titers than the corresponding shake flask process, likely due to improved pH control during cultivation. At the same time, reactor cultivation resulted in altered host cell metabolism, presumably caused by differences in shear stress and controlled process conditions. Glucose consumption rates increased approximately two-fold compared to shake flask cultivation, which is in agreement with previous studies and may reflect shear stress-associated metabolic adaptation (Gobel et al., 2023, Shen et al., 2024, Silva et al., 2023). In parallel, lactate concentrations at TOI and TOH were substantially elevated in the stirred tank reactors, reaching levels comparable to other fed-batch virus production processes using HEK293 cells (Silva et al., 2023, Shen et al., 2024), potentially due to incomplete glucose oxidation (Sidorenko et al., 2008). In contrast, amino acid consumption increased only moderately, indicating that no further adaptation of the feeding strategy was required after scale transfer. Interestingly, ammonium concentrations decreased during reactor cultivation and reached lower levels at harvest compared to shake flask production, potentially due to enhanced oxygen transfer and stripping effects, as similarly observed during perfusion-based influenza virus production in MDCK cells (Hein et al., 2021b).

Process evaluation demonstrated that fed-batch cultivation substantially improved influenza virus and OP7 chimera DIP titer compared to the reference batch process. In stirred tank reactors, fed-batch cultivation resulted in approximately six-fold higher infectious virus titers and a four-fold increase in VVP, highlighting the effectiveness of the feeding strategy in supporting productive infection at elevated cell densities. Interestingly, CSVYs remained largely comparable between fed-batch and batch cultivations, suggesting that the improved overall productivity was primarily driven by increased viable cell concentrations rather than enhanced per-cell productivity. Similar improvements in VVP following fed-batch optimization have also been reported for influenza virus and vesicular stomatitis virus production using HEK cells (Silva et al., 2023, Silva et al., 2025, Elahi et al., 2019). Importantly, the fed-batch process could also be successfully applied to OP7 chimera DIP production, where increased virus titers were accompanied by significantly enhanced biological interfering efficacy. HA-based virus titers reached 3.8 log_10_ HAU/100 µL, exceeding those of the batch reference process and approaching previously reported perfusion-based productions (Pelz et al., 2024). Notably, despite lower overall titers compared to perfusion cultivation, the biological interfering efficacy of fed-batch-derived material was comparable to perfusion-produced OP7 chimera DIPs.

Further process improvements during upstream processing could be achieved by using a high-density cryo cell bank to increase process economics by shortening the overall seed train time and by implementing an N-1 perfusion step to reduce the number of seed train steps (Zinnecker et al., 2025, Müller et al., 2022). In addition to seed train intensification, these approaches would also contribute to a leaner process and reduce both the risk of contamination and media consumption. In addition to seed train intensification, these approaches would also help to streamline the process and reduce both the risk of contamination and media consumption. As this would not require any changes to the final production bioreactor, and as perfusion wave bags with an integrated cell retention membrane are commercially available for N-1-perfusion, its implementation into the existing process would be easily achievable.

Previously, IAV DIPs were purified using membrane-based steric exclusion chromatography (Marichal-Gallardo et al., 2017), which proved to be a suitable method for preserving biological activity (Pelz et al., 2024, Hein et al., 2021a). With an additional focus on scalability, commercial availability and widespread industrial use, other chromatographic purification methods, such as modern pseudo-affinity chromatography using sulfated cellulose membrane adsorber for virus capture (Opitz et al., 2009, Carvalho et al., 2018) could be investigated in future. In addition, the formulation and virus stability have yet to be investigated.

Overall, the present study establishes fed-batch cultivation as a highly promising and industrially accessible production platform for antiviral OP7 chimera DIPs. In parallel, the generation of GMP-compliant MDCK-PB2(sus) cell banks and OP7 chimera seed virus banks provides an important basis for future GMP manufacturing for OP7 clinical evaluation. Together, these advances represent a significant step towards the clinical translation and large-scale manufacturing of OP7 chimera DIPs as a novel broad-spectrum antiviral treatment modality.

## Supplementary material

**Figure S1:**
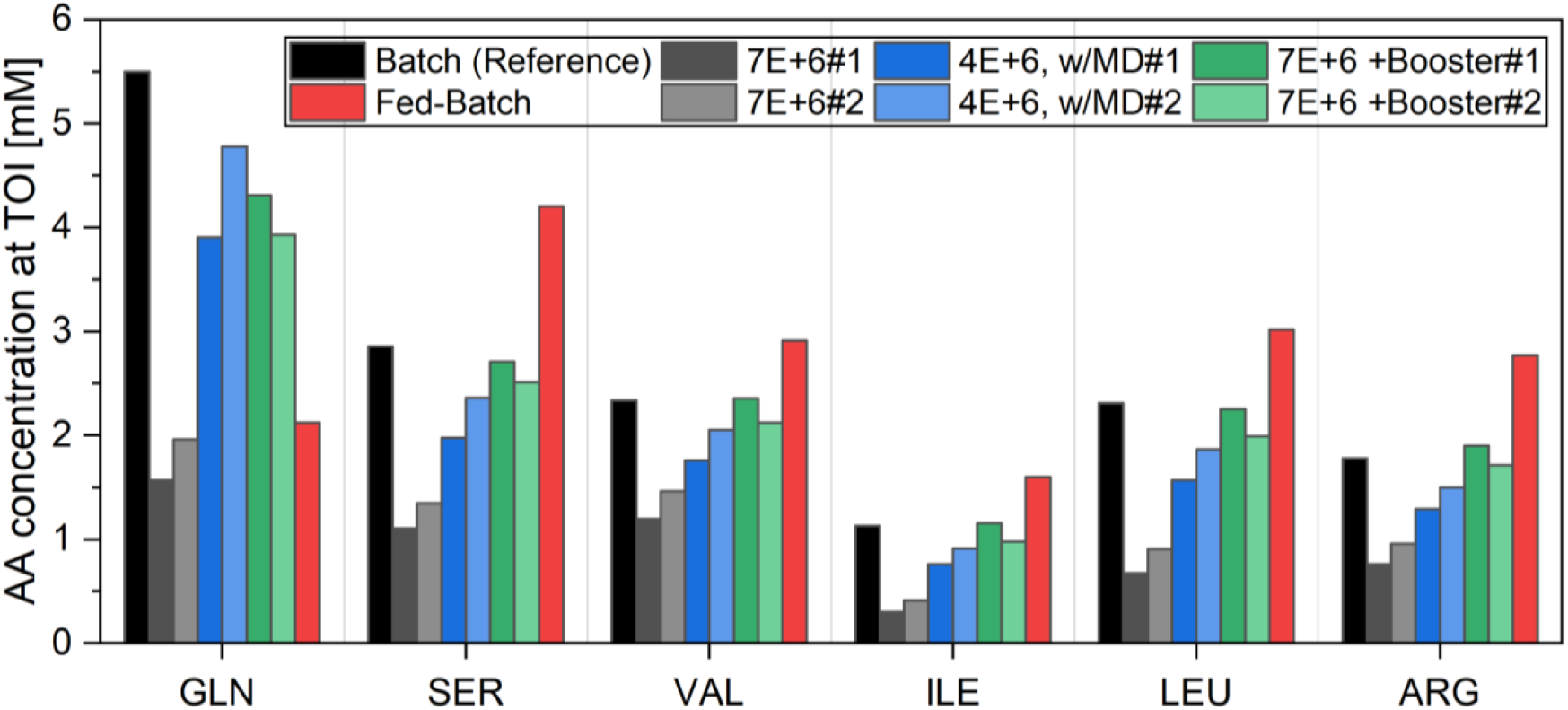
Amino acid (AA) concentrations at time of infection (TOI) during establishment of a fed-batch process for IAV production. The six most consumed amino acids glutamine (GLN), serine (SER), valine (VAL), isoleucine (ILE), leucine (LEU) and arginine (ARG) are shown. Note that the calibrated measurement range for the diluted samples (1:20 dilution) was between 1 and 8 mM. For fed-batch production, glutamine supplementation was substituted by GlutaMAX (L-alanyl-L-glutamine dipeptide) addition.

**Figure S2:**
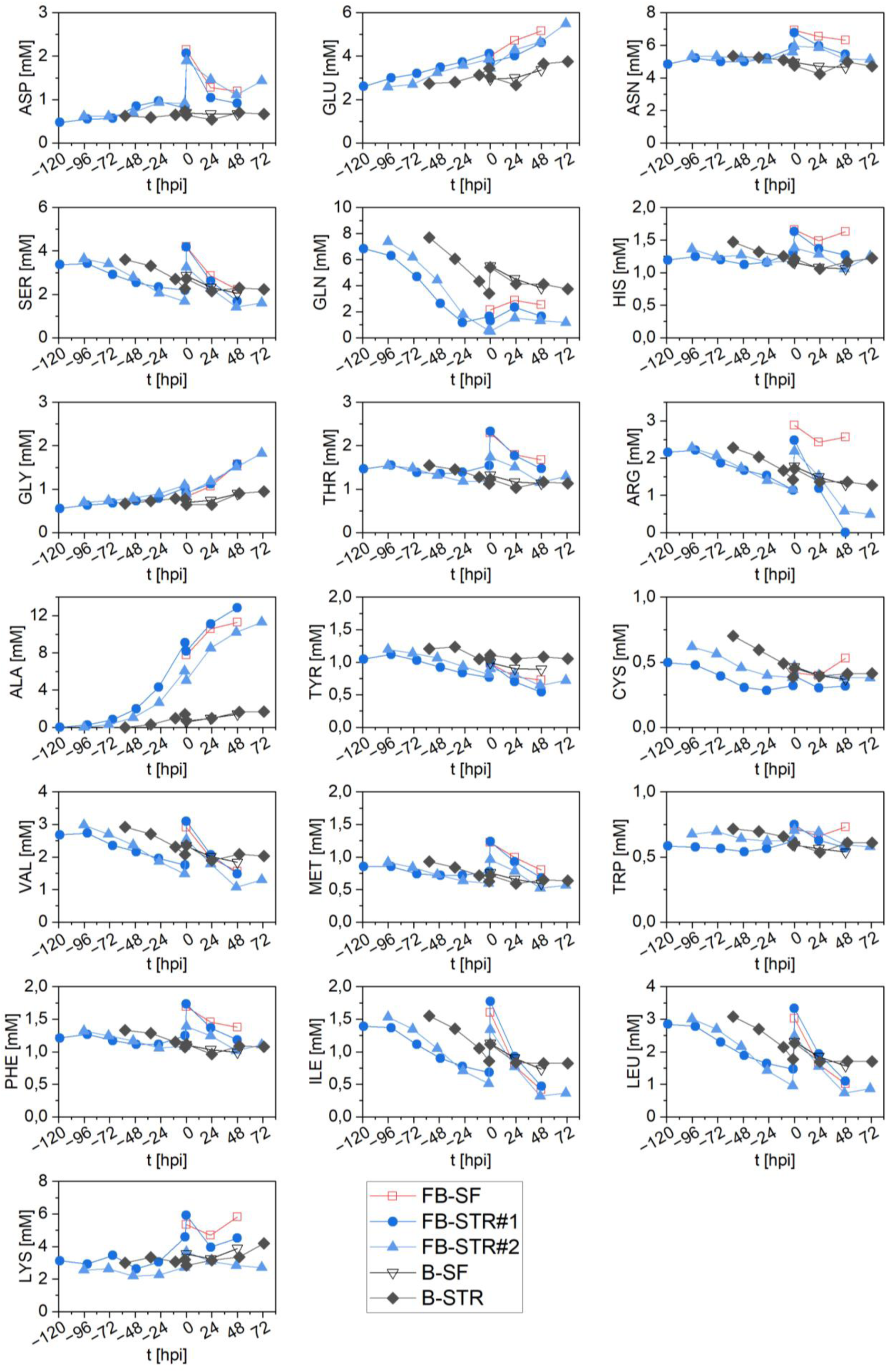
Amino acid concentrations during IAV-PR8 production in fed-batch (FB) and batch (B) mode in a 1L-STR compared to shake flask production (SF). Note that the calibrated measurement range for the diluted samples (1:20 dilution) was between 1 and 8 mM and that no proline data could be provided.

## References

1. Agu, I. & Diaz-Munoz, S. L. 2025. Host cell state: an overlooked factor impacting the production of influenza A deletion-containing viral genomes and non-infectious particles. mBio, 16, e0199125.

2. Alnaji, F. G. & Brooke, C. B. 2020. Influenza virus DI particles: Defective interfering or delightfully interesting? PLos Pathog, 16, e1008436.

3. Bdeir, N., Arora, P., Gartner, S., Hoffmann, M., Reichl, U., Pohlmann, S. & Winkler, M. 2019. A system for production of defective interfering particles in the absence of infectious influenza A virus. PLoS One, 14, e0212757.

4. Bissinger, T., Wu, Y., Marichal-Gallardo, P., Riedel, D., Liu, X., Genzel, Y., Tan, W. S. & Reichl, U. 2021. Towards integrated production of an influenza A vaccine candidate with MDCK suspension cells. Biotechnol Bioeng, 118, 3996–4013.

5. Carvalho, S. B., Fortuna, A. R., Wolff, M. W., Peixoto, C. M Alves, P., Reichl, U. & Carrondo, M. J. T. 2018. Purification of influenza virus-like PARTICLES using sulfated cellulose membrane adsorbers. Journal of Chemical Technology and Biotechnology, 93, 1988–1996.

6. Chaturvedi, S., Vasen, G., Pablo, M., Chen, X., Beutler, N., Kumar, A., Tanner, E., Illouz, S., Rahgoshay, D., Burnett, J., Holguin, L., Chen, P. Y., Ndjamen, B., Ott, M., Rodick, R., Rogers, T., Smith, D. M. & Weinberger, L. S. 2021. Identification of a therapeutic interfering particle-A single-dose SARS-CoV-2 antiviral intervention with a high barrier to resistance. Cell, 184, 6022–6036 e18.

7. Chen, C., Wong, H. E. & Goudar, C. T. 2018. Upstream process intensification and continuous manufacturing. Current Opinion in Chemical Engineering, 22, 191–198.

8. Dill, V., Ehret, J., Zimmer, A., Beer, M. & Eschbaumer, M. 2019. Cell Density Effects in Different Cell Culture Media and Their Impact on the Propagation of Foot-And-Mouth Disease Virus. Viruses-Basel, 11.

9. Dimmock, N. J. & Easton, A. J. 2015. Cloned Defective Interfering Influenza RNA and a Possible Pan-Specific Treatment of Respiratory Virus Diseases. Viruses, 7, 3768–88.

10. Dimmock, N. J., Rainsford, E. W., Scott, P. D. & Marriott, A. C. 2008. Influenza virus protecting Rna: an effective prophylactic and therapeutic antiviral. J Virol, 82, 8570–8.

11. Dogra, T., Pelz, L., Boehme, J. D., Kuechler, J., Kershaw, O., Marichal-Gallardo, P., Baelkner, M., Hein, M. D., Gruber, A. D., Benndorf, D., Genzel, Y., Bruder, D., Kupke, S. Y. & Reichl, U. 2023. Generation of “OP7 chimera” defective interfering influenza A particle preparations free of infectious virus that show antiviral efficacy in mice. Sci Rep, 13, 20936.

12. Easton, A. J., Scott, P. D., Edworthy, N. L., Meng, B., Marriott, A. C. & Dimmock, N. J. 2011. A novel broad-spectrum treatment for respiratory virus infections: influenza-based defective interfering virus provides protection against pneumovirus infection in vivo. Vaccine, 29, 2777–84.

13. Elahi, S. M., Shen, C. F. & Gilbert, R. 2019. Optimization of production of vesicular stomatitis virus (VSV) in suspension serum-free culture medium at high cell density. J Biotechnol, 289, 144–149.

14. Ferreira, T. B., Carrondo, M. J. & Alves, P. M. 2007. Effect of ammonia production on intracellular pH: Consequent effect on adenovirus vector production. J Biotechnol, 129, 433–8.

15. Genzel, Y., Ritter, J. B., Konig, S., Alt, R. & Reichl, U. 2005. Substitution of glutamine by pyruvate to reduce ammonia formation and growth inhibition of mammalian cells. Biotechnol Prog, 21, 58–69.

16. Gobel, S., Jaen, K. E., Dorn, M., Neumeyer, V., Jordan, I., Sandig, V., Reichl, U., Altomonte, J. & Genzel, Y. 2023. Process intensification strategies toward cell culture-based high-yield production of a fusogenic oncolytic virus. Biotechnol Bioeng, 120, 2639–2657.

17. Granicher, G., Coronel, J., Pralow, A., Marichal-Gallardo, P., Wolff, M., Rapp, E., Karlas, A., Sandig, V., Genzel, Y. & Reichl, U. 2019. Efficient influenza A virus production in high cell density using the novel porcine suspension cell line PBG.PK2.1. Vaccine, 37, 7019–7028.

18. Hein, M. D., Arora, P., Marichal-Gallardo, P., Winkler, M., Genzel, Y., Pohlmann, S., Schughart, K., Kupke, S. Y. & Reichl, U. 2021a. Cell culture-based production and in vivo characterization of purely clonal defective interfering influenza virus particles. BMC Biol, 19, 91.

19. Hein, M. D., Chawla, A., Cattaneo, M., Kupke, S. Y., Genzel, Y. & Reichl, U. 2021b. Cell culture-based production of defective interfering influenza A virus particles in perfusion mode using an alternating tangential flow filtration system. Appl Microbiol Biotechnol, 105, 7251–7264.

20. Hein, M. D., Kazenmaier, D., Van Heuvel, Y., Dogra, T., Cattaneo, M., Kupke, S. Y., Stitz, J., Genzel, Y. & Reichl, U. 2023. Production of retroviral vectors in continuous high cell density culture. Appl Microbiol Biotechnol, 107, 5947–5961.

21. Hein, M. D., Kollmus, H., Marichal-Gallardo, P., Puttker, S., Benndorf, D., Genzel, Y., Schughart, K., Kupke, S. Y. & Reichl, U. 2021c. OP7, a novel influenza A virus defective interfering particle: production, purification, and animal experiments demonstrating antiviral potential. Appl Microbiol Biotechnol, 105, 129–146.

22. Henry, O., Dormond, E., Perrier, M. & Kamen, A. 2004. Insights into adenoviral vector production kinetics in acoustic filter-based perf usion cultures. Biotechnology and Bioengineering, 86, 765–774.

23. Holmes, E. C., Hurt, A. C., Dobbie, Z., Clinch, B., Oxford, J. S. & Piedra, P. A. 2021. Understanding the Impact of Resistance to Influenza Antivirals. Clinical Microbiology Reviews, 34.

24. Joe, C. C. D., Jiang, J., Linke, T., Li, Y., Fedosyuk, S., Gupta, G., Berg, A., Segireddy, R. R., Mainwaring, D., Joshi, A., Cashen, P., Rees, B., Chopra, N., Nestola, P., Humphreys, J., Davies, S., Smith, N., Bruce, S., Verbart, D., Bormans, D., Knevelman, C., Woodyer, M., Davies, L., Cooper, L., Kapanidou, M., Bleckwenn, N., Pappas, D., Lambe, T., Smith, D. C., Green, C. M., Venkat, R., Ritchie, A. J., Gilbert, S. C., Turner, R. & Douglas, A. D. 2022. Manufacturing a chimpanzee adenovirus-vectored SARS-CoV-2 vaccine to meet global needs. Biotechnol Bioeng, 119, 48–58.

25. Jones, J. C., Yen, H. L., Adams, P., Armstrong, K. & Govorkova, E. A. 2023. Influenza antivirals and their role in pandemic preparedness. Antiviral Research, 210.

26. Kalamvoki, M. & Norris, V. 2022. A Defective Viral Particle Approach to COVID-19. Cells, 11.

27. Kalbfuss, B., Knochlein, A., Krober, T. & Reichl, U. 2008. Monitoring influenza virus content in vaccine production: precise assays for the quantitation of hemagglutination and neuraminidase activity. Biologicals, 36, 145–61.

28. Kegel, N. B., Kaufmann, A., Matrosovich, M., Bauer, S. & Dorna, J. 2025. Decomposition of L-glutamine and accumulation of ammonium in cell culture media inhibit infectivity of influenza viruses. Virology, 602.

29. Killip, M. J., Fodor, E. & Randall, R. E. 2015. Influenza virus activation of the interferon system. Virus Res, 209, 11–22.

30. Krammer, F., Smith, G. J. D., Fouchier, R. A. M., Peiris, M., Kedzierska, K., Doherty, P. C., Palese, P., Shaw, M. L., Treanor, J., Webster, R. G. & Garcia-Sastre, A. 2018. Influenza. Nat Rev Dis Primers, 4, 3.

31. Kuchler, J., Puttker, S., Lahmann, P., Genzel, Y., Kupke, S., Benndorf, D. & Reichl, U. 2022. Absolute quantification of viral proteins during single-round replication of MDCK suspension cells. J Proteomics, 259, 104544.

32. Kupke, S. Y., Riedel, D., Frensing, T., Zmora, P. & Reichl, U. 2019. A Novel Type of Influenza A Virus-Derived Defective Interfering Particle with Nucleotide Substitutions in Its Genome. J Virol, 93.

33. Lavado-Garcia, J., Perez-Rubio, P., Cervera, L. & Godia, F. 2022. The cell density effect in animal cell-based bioprocessing: Questions, insights and perspectives. Biotechnology Advances, 60.

34. Levi, L. I., Rezelj, V. V., Henrion-Lacritick, A., Erazo, D., Boussier, J., Vallet, T., Bernhauerova, V., Suzuki, Y., Carrau, L., Weger-Lucarelli, J., Saleh, M. C. & Vignuzzi, M. 2021. Defective viral genomes from chikungunya virus are broad-spectrum antivirals and prevent virus dissemination in mosquitoes. PLoS Pathog, 17, e1009110.

35. Li, D., Lin, M. H., Rawle, D. J., Jin, H., Wu, Z., Wang, L., Lor, M., Hussain, M., Aaskov, J. & Harrich, D. 2021. Dengue virus-free defective interfering particles have potent and broad anti-dengue virus activity. Commun Biol, 4, 557.

36. Lin, M. H., Li, D., Tang, B., Li, L., Suhrbier, A. & Harrich, D. 2022. Defective Interfering Particles with Broad-Acting Antiviral Activity for Dengue, Zika, Yellow Fever, Respiratory Syncytial and SARS-CoV-2 Virus Infection. Microbiol Spectr, 10, e0394922.

37. Lohmann, J. J. G., Le, M., Alnaji, F. G., Zolotareva, O., Baumbach, J. & Laske, T. 2025. Meta-analysis of genomic characteristics for antiviral influenza defective interfering particle prioritization. NAR Genom Bioinform, 7, lqaf031.

38. Marichal-Gallardo, P., Pieler, M. M., Wolff, M. W. & Reichl, U. 2017. Steric exclusion chromatography for purification of cell culture-derived influenza A virus using regenerated cellulose membranes and polyethylene glycol. J Chromatogr A, 1483, 110–119.

39. Matanguihan, C. & Wu, P. 2022. Upstream continuous processing: recent advances in production of biopharmaceuticals and challenges in manufacturing. Curr Opin Biotechnol, 78, 102828.

40. Mendes, M. & Russell, A. B. 2021. Library-based analysis reveals segment and length dependent characteristics of defective influenza genomes. PLoS Pathog, 17, e1010125.

41. Müller, J., Ott, V., Eibl, D. & Eibl, R. 2022. Seed Train Intensification Using an Ultra-High Cell Density Cell Banking Process. Processes, 10.

42. Nypaver, C., Dehlinger, C. & Carter, C. 2021. Influenza and Influenza Vaccine: A Review. J Midwifery Womens Health, 66, 45–53.

43. Opitz, L., Lehmann, S., Reichl, U. & Wolff, M. W. 2009. Sulfated Membrane Adsorbers for Economic Pseudo-Affinity Capture of Influenza Virus Particles. Biotechnology and Bioengineering, 103, 1144–1154.

44. Pelz, L., Dogra, T., Marichal-Gallardo, P., Hein, M. D., Hemissi, G., Kupke, S. Y., Genzel, Y. & Reichl, U. 2024. Production of antiviral “OP7 chimera” defective interfering particles free of infectious virus. Appl Microbiol Biotechnol, 108, 97.

45. Pelz, L., Piagnani, E., Marsall, P., Wynserski, N., Hein, M. D., Marichal-Gallardo, P., Kupke, S. Y. & Reichl, U. 2023. Broad-Spectrum Antiviral Activity of Influenza A Defective Interfering Particles against Respiratory Syncytial, Yellow Fever, and Zika Virus Replication In Vitro. Viruses, 15.

46. Pitchai, F. N. N., Tanner, E. J., Khetan, N., Vasen, G., Levrel, C., Kumar, A. J., Pandey, S., Ordonez, T., Barnette, P., Spencer, D., Jung, S. Y., Glazier, J., Thompson, C., Harvey-Vera, A., Son, H. I., Son, H. I., Strathdee, S. A., Holguin, L., Urak, R., Burnett, J., Burgess, W., Busman-Sahay, K., Estes, J. D., Hessell, A., Fennessey, C. M., Keele, B. F., Haigwood, N. L. & Weinberger, L. S. 2024. Engineered deletions of HIV replicate conditionally to reduce disease in nonhuman primates. Science, 385, eadn5866.

47. Rand, U., Kupke, S. Y., Shkarlet, H., Hein, M. D., Hirsch, T., Marichal-Gallardo, P., Cicin-Sain, L., Reichl, U. & Bruder, D. 2021. Antiviral Activity of Influenza A Virus Defective Interfering Particles against SARS-CoV-2 Replication In Vitro through Stimulation of Innate Immunity. Cells, 10.

48. Ranum, J. N., Ledwith, M. P., Alnaji, F. G., Diefenbacher, M., Orton, R., Sloan, E., Guereca, M., Feltman, E. M., Smollett, K., Da Silva Filipe, A., Conley, M., Russell, A. B., Brooke, C. B., Hutchinson, E. & Mehle, A. 2024. Cryptic proteins translated from deletion-containing viral genomes dramatically expand the influenza virus proteome. Nucleic Acids Res, 52, 3199–3212.

49. Rast, L. I., Rouzine, I. M., Rozhnova, G., Bishop, L., Weinberger, A. D. & Weinberger, L. S. 2016. Conflicting Selection Pressures Will Constrain Viral Escape from Interfering Particles: Principles for Designing Resistance-Proof Antivirals. PLoS Comput Biol, 12, e1004799.

50. Rezelj, V. V., Carrau, L., Merwaiss, F., Levi, L. I., Erazo, D., Tran, Q. D., Henrion-Lacritick, A., Gausson, V., Suzuki, Y., Shengjuler, D., Meyer, B., Vallet, T., Weger-Lucarelli, J., Bernhauerova, V., Titievsky, A., Sharov, V., Pietropaoli, S., Diaz-Salinas, M. A., Legros, V., Pardigon, N., Barba-Spaeth, G., Brodsky, L., Saleh, M. C. & Vignuzzi, M. 2021. Defective viral genomes as therapeutic interfering particles against flavivirus infection in mammalian and mosquito hosts. Nat Commun, 12, 2290.

51. Rezelj, V. V., Levi, L. I. & Vignuzzi, M. 2018. The defective component of viral populations. Curr Opin Virol, 33, 74–80.

52. Rüdiger, D., Piasecka, J., Kuchler, J., Pontes, C., Laske, T., Kupke, S. Y. & Reichl, U. 2024. Mathematical model calibrated to in vitro data predicts mechanisms of antiviral action of the influenza defective interfering particle “OP7”. iScience, 27, 109421.

53. Salvi, A., Quillan, J. M. & Sadee, W. 2002. Monitoring intracellular ph changes in response to osmotic stress and membrane transport activity using 5-chloromethylfluorescein. Aaps PharmSci, 4, E21.

54. Scott, P. D., Meng, B., Marriott, A. C., Easton, A. J. & Dimmock, N. J. 2011. Defective interfering influenza A virus protects in vivo against disease caused by a heterologous influenza B virus. J Gen Virol, 92, 2122–2132.

55. Sharov, V., Rezelj, V. V., Galatenko, V. V., Titievsky, A., Panov, J., Chumakov, K., Andino, R., Vignuzzi, M. & Brodsky, L. 2021. Intra- and Inter-cellular Modeling of Dynamic Interaction between Zika Virus and Its Naturally Occurring Defective Viral Genomes. J Virol, 95, e0097721.

56. Shen, C. F., Rodenbrock, A., Lanthier, S., Burney, E. & Loignon, M. 2024. Optimization of Culture Media and Feeding Strategy for High Titer Production of an Adenoviral Vector in HEK 293 Fed-Batch Culture. Vaccines (Basel), 12.

57. Sidorenko, Y., Schulze-Horsel, J., Voigt, A., Reichl, U. & Kienle, A. 2008. Stochastic population balance modeling of influenza virus replication in vaccine production processes. Chemical Engineering Science, 63, 157–169.

58. Silva, C. A. T., Kamen, A. A. & Henry, O. 2021. Recent advances and current challenges in process intensification of cell culture-based influenza virus vaccine manufacturing. Canadian Journal of Chemical Engineering, 99, 2525–2535.

59. Silva, C. A. T., Kamen, A. A. & Henry, O. 2023. Intensified Influenza Virus Production in Suspension HEK293SF Cell Cultures Operated in Fed-Batch or Perfusion with Continuous Harvest. Vaccines (Basel), 11.

60. Silva, C. A. T., Kamen, A. A. & Henry, O. 2025. Fed-batch strategies for intensified rVSV vector production in high cell density cultures of suspension HEK293 cells. Biotechnol Prog, 41, e3506.

61. Smither, S. J., Garcia-Dorival, I., Eastaugh, L., Findlay, J. S., O’brien, L. M., Carruthers, J., Williamson, E. D., Molina-Paris, C., Hiscox, J. A. & Laws, T. R. 2020. An Investigation of the Effect of Transfected Defective, Ebola Virus Genomes on Ebola Replication. Front Cell Infect Microbiol, 10, 159.

62. Tanner, A. R., Dorey, R. B., Brendish, N. J. & Clark, T. W. 2021. Influenza vaccination: protecting the most vulnerable. Eur Respir Rev, 30.

63. Tapia, K., Kim, W. K., Sun, Y., Mercado-Lopez, X., Dunay, E., Wise, M., Adu, M. & Lopez, C. B. 2013. Defective viral genomes arising in vivo provide critical danger signals for the triggering of lung antiviral immunity. PLos Pathog, 9, e1003703.

64. Vazquez-Ramirez, D., Jordan, I., Sandig, V., Genzel, Y. & Reichl, U. 2019. High titer MVA and influenza A virus production using a hybrid fed-batch/perfusion strategy with an ATF system. Appl Microbiol Biotechnol, 103, 3025–3035.

65. Vignuzzi, M. & Lopez, C. B. 2019. Defective viral genomes are key drivers of the virus-host interaction. Nat Microbiol, 4, 1075–1087.

66. Welch, S. R., Spengler, J. R., Harmon, J. R., Coleman-Mccray, J. D., Scholte, F. E. M., Genzer, S. C., Lo, M. K., Montgomery, J. M., Nichol, S. T. & Spiropoulou, C. F. 2022. Defective Interfering Viral Particle Treatment Reduces Clinical Signs and Protects Hamsters from Lethal Nipah Virus Disease. mBio, 13, e0329421.

67. Wu, Y., Bissinger, T., Genzel, Y., Liu, X., Reichl, U. & Tan, W. S. 2021. High cell density perfusion process for high yield of influenza A virus production using MDCK suspension cells. Appl Microbiol Biotechnol, 105, 1421–1434.

68. Xiao, Y., Lidsky, P. V., Shirogane, Y., Aviner, R., Wu, C. T., Li, W., Zheng, W., Talbot, D., Catching, A., Doitsh, G., Su, W., Gekko, C. E., Nayak, A., Ernst, J. D., Brodsky, L., Brodsky, E., Rousseau, E., Capponi, S., Bianco, S., Nakamura, R., Jackson, P. K., Frydman, J. & Andino, R. 2021. A defective viral genome strategy elicits broad protective immunity against respiratory viruses. Cell, 184, 6037–6051 e14.

69. Yin, J. 2025. Matryoshka Dolls of Virus Evolution: the Internal Forces That Shape Viral Fate. Integr Comp Biol.

70. Yount, J. S., Gitlin, L., Moran, T. M. & Lopez, C. B. 2008. MDA5 participates in the detection of paramyxovirus infection and is essential for the early activation of dendritic cells in response to Sendai Virus defective interfering particles. J Immunol, 180, 4910–8.

71. Zhao, H., Zhang, C., Lam, H., Meng, X., Peng, Z., Yeung, M. L., Chan, J. F., Kai-Wang To, K. & Yuen, K. Y. 2022. Peptidic defective interfering gene nanoparticles against Omicron, Delta SARS-CoV-2 variants and influenza A virus in vivo. Signal Transduct Target Ther, 7, 266.

72. Zinnecker, T., Wicke, E., Reichl, U., Göbel, S. & Genzel, Y. 2025. Seed Train Intensification and TFDF-Based Perfusion for MDCK Cell-Based Influenza A Virus Production. Processes, 13.

